## Supplemental Fig. 1-7 & Table 1-2 for "Double NPY motifs at the N-terminus of Sso2 synergistically bind Sec3 to promote membrane fusion"

#### Legends for supplemental figures

**Fig. S1. Constructs and purification of the Sso2/Sec3 complex.** (**A**) Schematics of the constructs of Sso2 (aa1–270) and Sec3 (aa75–260) used in our studies. (**B**) Size-exclusion chromatography elution profiles of Sso2, Sec3, and a mixture of Sso2 and Sec3 (molar ratio 1:3). The SDS PAGE gel shows proteins in the mixture ("Load") and fractions from the two elution peaks of the mixture. The first peak contains the Sso2/Sec3 complex with both proteins; the second one contains only excessive Sec3 alone.

**Fig. S2. Structure of the Sso2/Sec3 complex and conservation analysis.** (**A**) Two different representations of the Sso2/Sec3 complex structure show the two separate interaction sites between the two proteins. The NPY motif of Sso2 is shown as magenta sticks. Dash lines depict the regions that are invisible in the crystal structure. (**B**) Sequence alignment of multiple Sso2 homologs. Highly and relatively conserved residues are shaded in dark and light blue, respectively. Secondary structures are shown above the aligned sequences. The two highly conserved NPY motifs at the N terminus of Sso2 are depicted as magenta blocks. Dash lines represent the unstructured loops that are invisible in the crystal structure. The C-terminal twenty-five residues of Sso2, including the hydrophobic transmembrane helix, is colored grey and was excluded from our protein expression construct.

**Fig. S3. Local chemical environment around the NPY motifs in the Sso2/Sec3 complex structure.** (**A**) Stereo view of the enlarged NPY binding sites of the two superimposed structures. Sec3 and the NPY motifs of Sso2 are shown as ribbons and sticks, respectively. Water molecules bound to the NPY motifs (directly or indirectly) are depicted as spheres with the same colors as the corresponding NPY motifs. (**B** & **C**) Separate stereo views of the two NPY binding sites. Sec3 molecules are shown as ribbons, with those residues hydrogen bonded to Sso2 or Sso2-bound water molecules additionally shown as sticks. The NPY motifs and waters are shown as sticks and spheres, respectively. All hydrogen bonds are depicted as dash lines. (**D** & **E**) Details of interactions between the NPY motifs of Sso2 (purple) and Sec3 (gray). The plots were generated using DIMPLOT in the LigPlot plus suite. Residues involved in hydrogen bond formation are shown as ball-and-stick, with oxygen, nitrogen, and carbon atoms colored in red, blue, and gray, respectively. Water molecules mediating intermolecular hydrogen bond formation are shown as cyan-colored spheres. Green dash lines indicate hydrogen bonds. Non-bonded residues involved in hydrophobic interactions are shown as spoked arcs. For clarity, only three of the seven bound water molecules are shown.

**Fig. S4. Growth effect and influence on Bgl2 and invertase secretion of s*so2* mutants of its first NPY motif.** (**A**) The sequence of *sso2* mutants generated by site-directed mutagenesis. Residues in the first NPY motif that were mutated to Ala (A) in all mutants (*i.e.* M1-M5) are shown in bold italics. (**B**) The *sso2* mutations in an *sso1Δ* background partially inhibit cells growth at 37$^o$C. Control *sso1Δ SSO2* or *sso1Δ sso2* mutant cells were grown overnight in YPD medium. An aliquot (0.2 $OD_{600}$ units) of cells from each strain was collected, serially diluted by 5 fold and spotted onto YPD plates. Plates were incubated at 25$^o$C, 34 $^o$C or 37$^o$C for 2 days. (**C**) Control *sso1Δ SSO2* strain and *sso1Δ sso2* mutants were grown as described in Figure 3. The external and internal Bgl2 pools were detected by western blotting. Several mutations in *SSO2* caused inhibition of Bgl2 secretion. (**D**) Quantitation of internal Bgl2. Results were analyzed based on four independent experiments. Error bar represents SD, n=5. *p<0.05. (**E**) Invertase secretion was determined in *sso2* mutants. Strains were grown as in Figure 3. Samples were processed and quantified as described in *Materials and Methods.* Three independent experiments were performed. Error bar represents SD, n=3. *p < 0.05. Invertase accumulation was calculated by the formula of $(Ext_{45m}-Ext_{0m}) /[(Ext_{45m}-Ext_{0m}) + (Int_{45m}-Int_{0m})]$.

**Fig. S5. *In vitro* binding assays to check the interaction between Sec3 and WT or mutant M7 of Sso2.** (**A**) Native gel electrophoresis shows that WT Sso2 (aa1-270) interacted with the Sec3 PH domain, with the slowly migrating complex bands becoming stronger with increasing amounts of Sec3. In contrast, the M7 mutant of Sso2 (aa1-270) shows no interaction. (**B**) The same set of samples in (A) were checked on an SDS PAGE gel to visualize the proteins. (**C**) Size exclusion chromatography elution profiles of either WT (blue) or mutant M7 (orange) Sso2 (aa1-270) mixed with the Sec3 PH domain. (**D**) The mixtures and samples from the elution peaks in (C) were checked on an SDS PAGE gel.

**Fig. S6. Co-immunoprecipitation of Sec3-3×Flag and Sso2 from yeast extract.** A control yeast strain expressing untagged Sec3 and Sso2 (NY3305) was grown in YPD and strains expressing Sec3-3×Flag and Sso2 (NY3447) or Sec3-3×Flag and Sso2-M7 (NY3448) were grown in SC-Leu medium at 25$^{o}$C overnight to an $OD_{600}$ around 1.0. Cell lysates were prepared as described in Methods section, and Sec3-3×Flag was immunoprecipitated from cell lysate with anti-Flag beads. Cell lysates and immunoprecipitated proteins (IPs) were blotted with anti-Flag antibody to detect Sec3-3×Flag (top panel) and anti-Sso antibody to detect Sso2 (bottom panel). The input was 1.7% of the lysate for Sec3-3×Flag and 0.25% for Sso2.

**Fig. S7. A hypothetical working model for how Sso2 recruits Sec3 in vesicle docking.** The NPY motifs at the N-terminus of Sso2 first reach out to dock into the complementary pocket on the Sec3 PH domain, which then pulls back and allows Sec3 to bind to the middle of the helical bundle of Sso2 (helices Ha, Hb and Hc are shown as semitransparent white cylinders, and H3 as an orange cylinder). Binding of Sec3 allosterically destabilizes the linker connecting Hc and H3 of Sso2 and primes downstream events in SNARE complex assembly.

Fig. S1

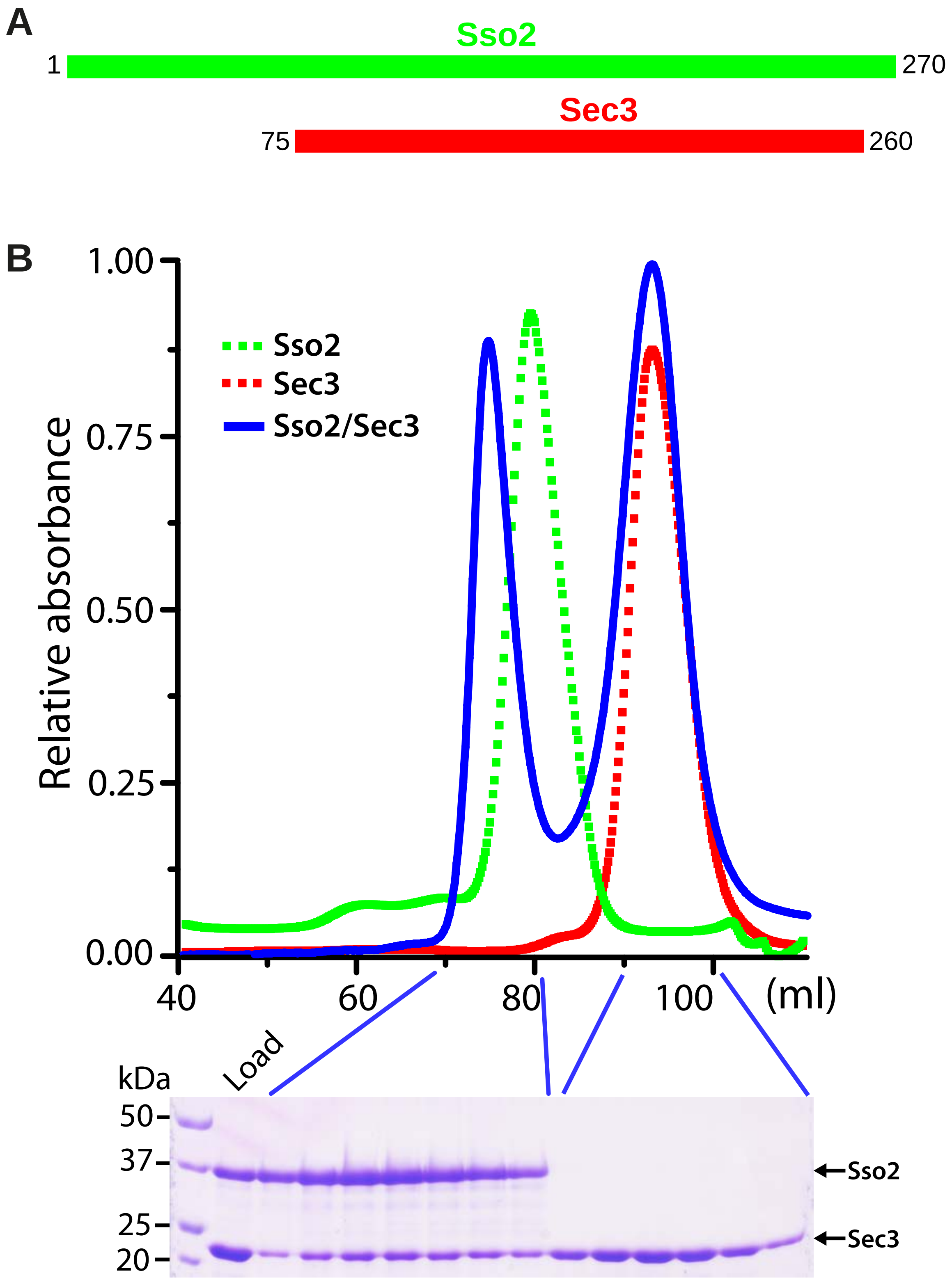
A
Sso2
1
270
Sec3
75
260
B
Relative absorbance
1.00
0.75
0.50
0.25
0.00
Sso2
Sec3
Sso2/Sec3
40
60
80
100
(ml)
Load
kDa
50
37
25
20
Sso2
Sec3

### Fig. S2

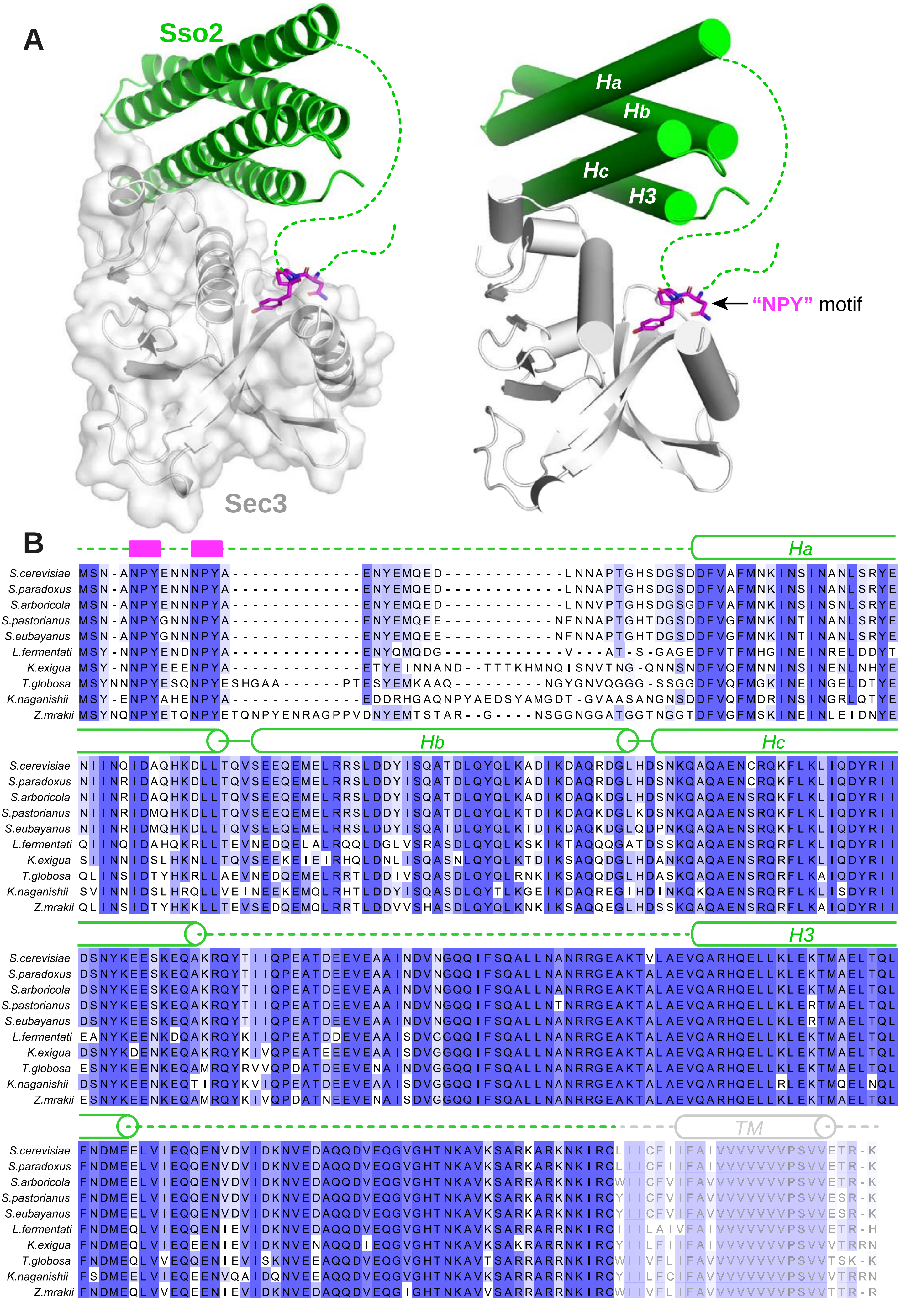
A
Sso2
Sec3
Ha
Hb
Hc
H3
"NPY" motif
B
Ha
Hb
Hc
H3
TM
S.cerevisiae
S.paradoxus
S.arboricola
S.pastorianus
S.eubayanus
L.fermentati
K.exigua
T.globosa
K.naganishii
Z.mrakii

Fig. S3

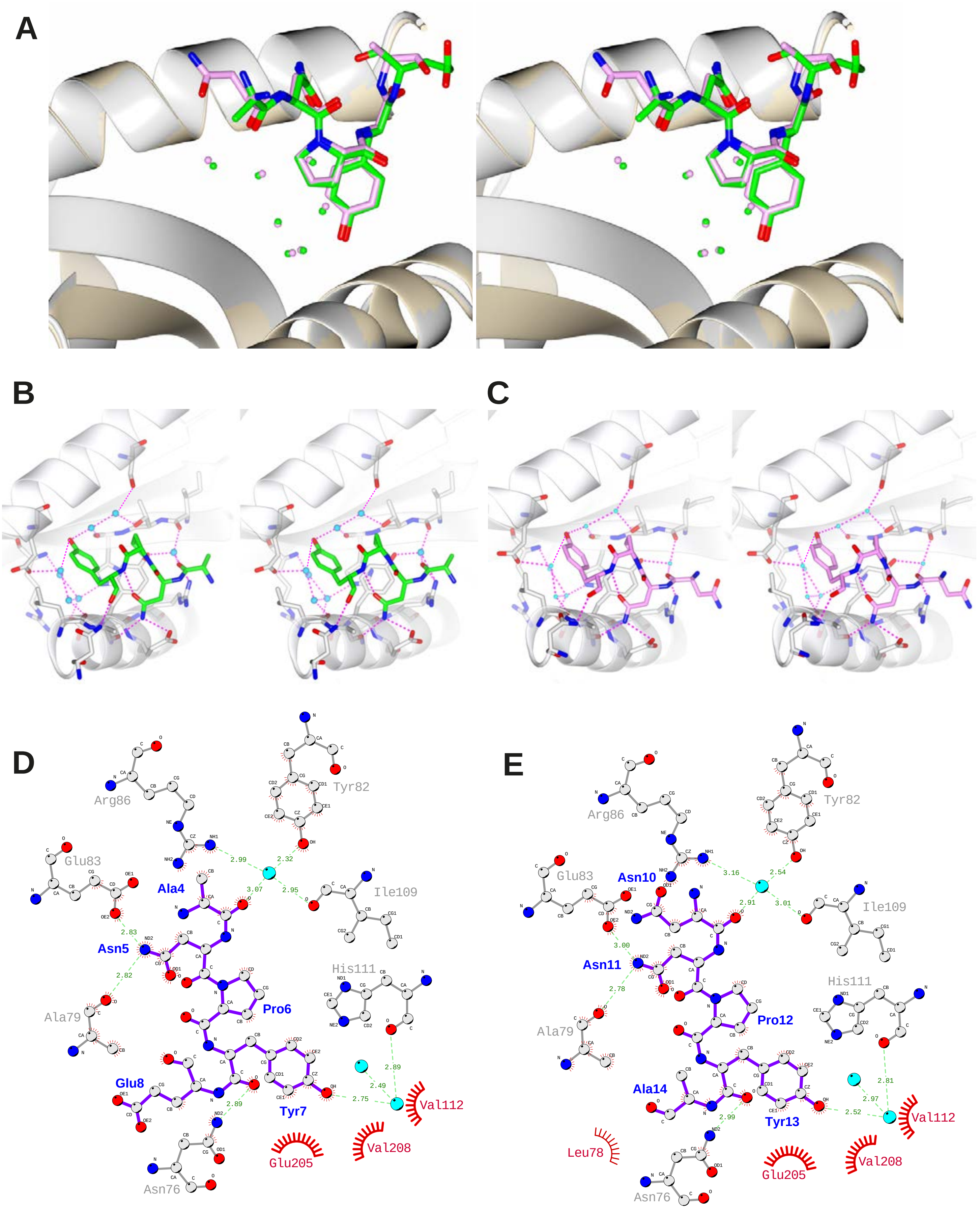
A
B
C
D
Arg86
Tyr82
Glu83
Ala4
Ile109
Asn5
His111
Pro6
Ala79
Glu8
Tyr7
Val112
Val208
Glu205
Asn76
E
Arg86
Tyr82
Glu83
Asn10
Ile109
Asn11
His111
Pro12
Ala79
Ala14
Tyr13
Val112
Leu78
Glu205
Val208
Asn76

### Fig. S4

**A** Sequences of Sso2 and *sso2* mutants (M1-M5):

| | 1 | 2 | 3 | 4 | 5 | 6 | 7 | 8 | 9 | 10 | 11 | 12 | 13 | 14 | 15 | 16 |
|---|---|---|---|---|---|---|---|---|---|---|---|---|---|---|---|---|
| *SSO2* | M | S | N | A | N | P | Y | E | N | N | N | P | Y | A | E | N |
| *M1* | M | S | N | A | N | P | Y | ***A*** | N | N | N | P | Y | A | E | N |
| *M2* | M | S | N | A | N | P | ***A*** | E | N | N | N | P | Y | A | E | N |
| *M3* | M | S | N | A | ***A*** | ***A*** | Y | E | N | N | N | P | Y | A | E | N |
| *M4* | M | S | N | A | ***A*** | ***A*** | ***A*** | E | N | N | N | P | Y | A | E | N |
| *M5* | M | S | N | A | ***A*** | ***A*** | ***A*** | ***A*** | N | N | N | P | Y | A | E | N |

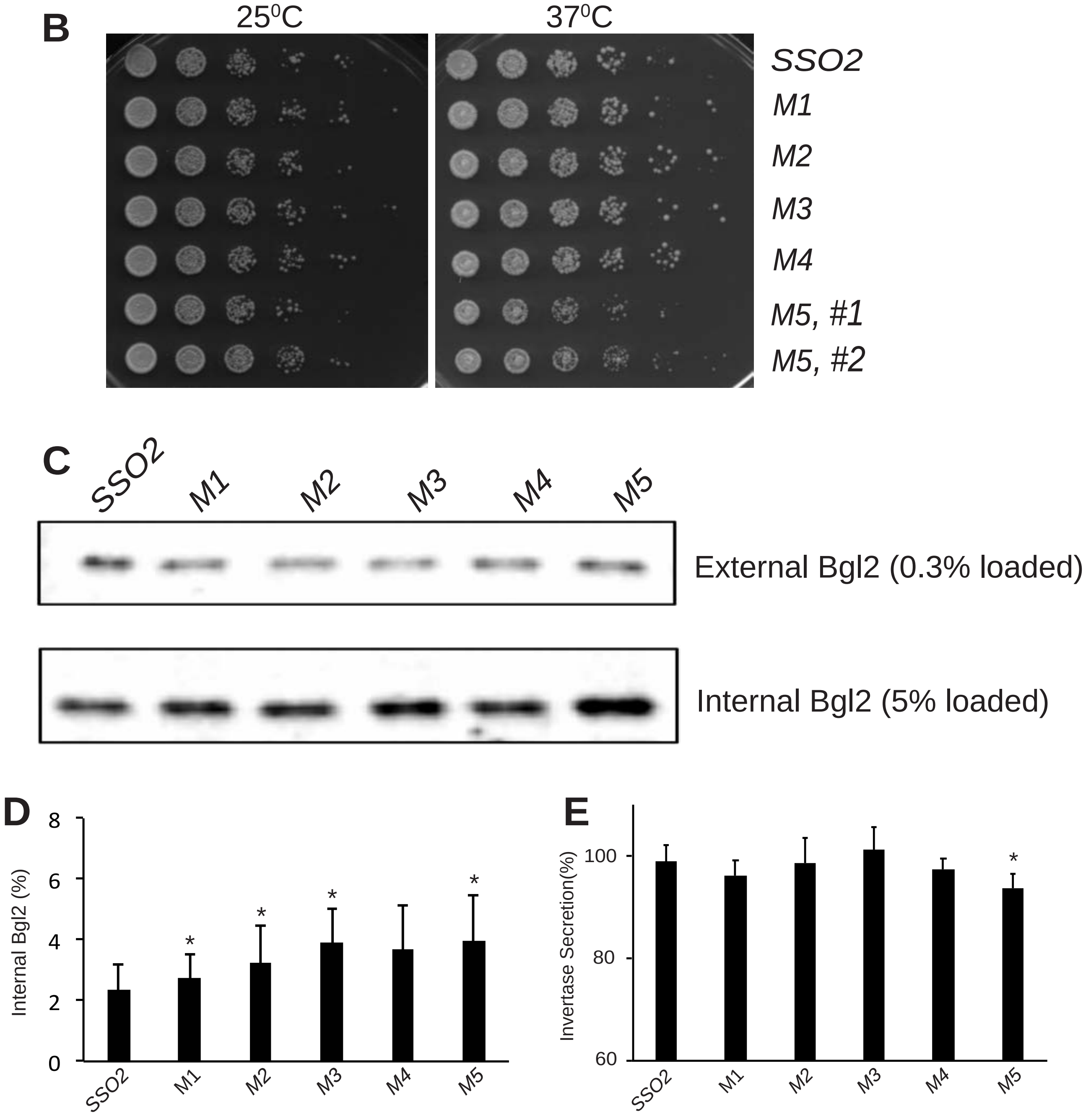

Fig. S5

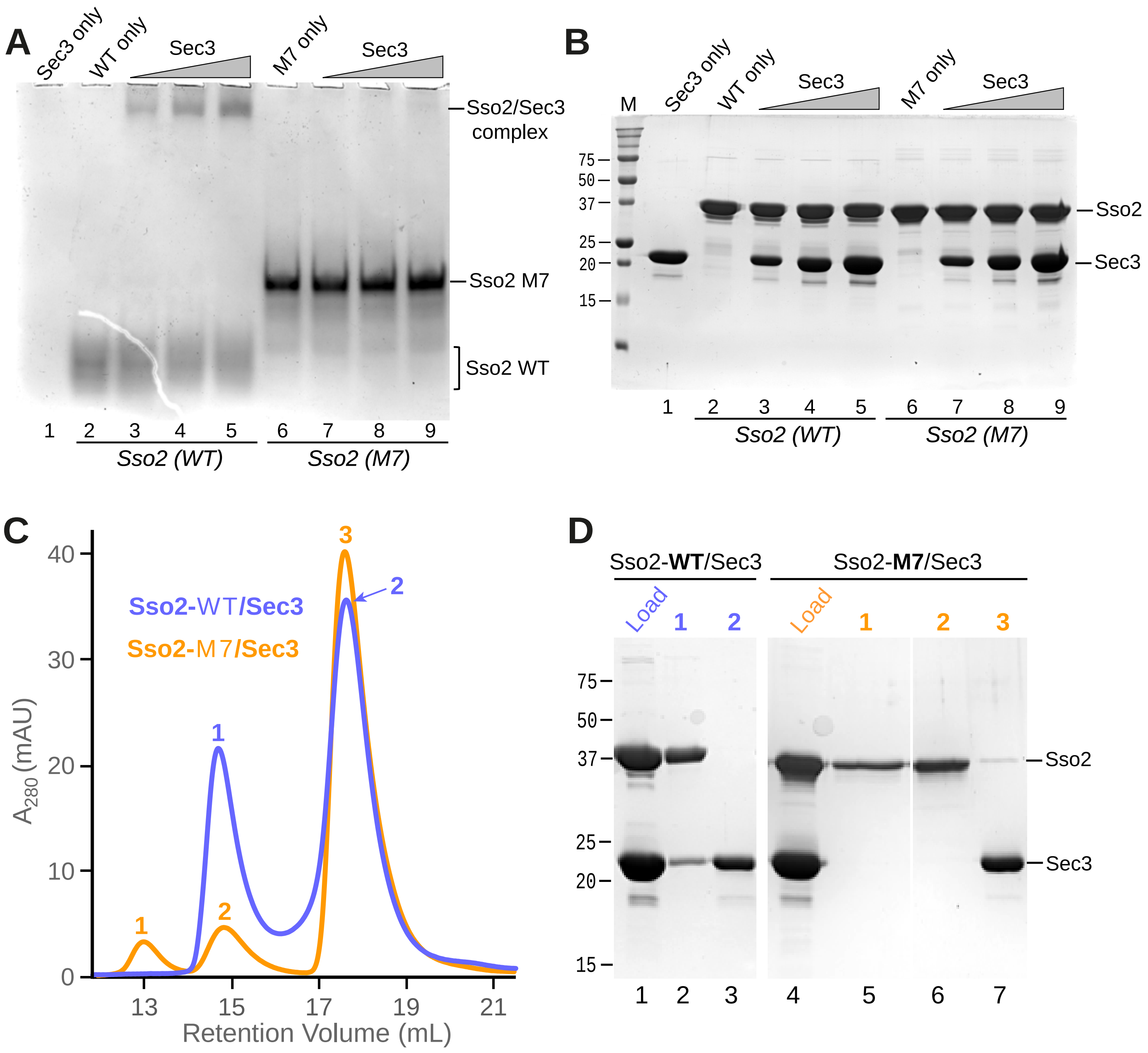
A
Sec3 only
WT only
Sec3
M7 only
Sec3
Sso2/Sec3 complex
Sso2 M7
Sso2 WT
1 2 3 4 5 6 7 8 9
Sso2 (WT)
Sso2 (M7)
B
M
Sec3 only
WT only
Sec3
M7 only
Sec3
75
50
37
25
20
15
Sso2
Sec3
1 2 3 4 5 6 7 8 9
Sso2 (WT)
Sso2 (M7)
C
Sso2-WT/Sec3
Sso2-M7/Sec3
40
30
20
10
0
A280 (mAU)
13 15 17 19 21
Retention Volume (mL)
D
Sso2-WT/Sec3
Sso2-M7/Sec3
Load 1 2
Load 1 2 3
75
50
37
25
20
15
Sso2
Sec3
1 2 3 4 5 6 7

**Fig. S6**

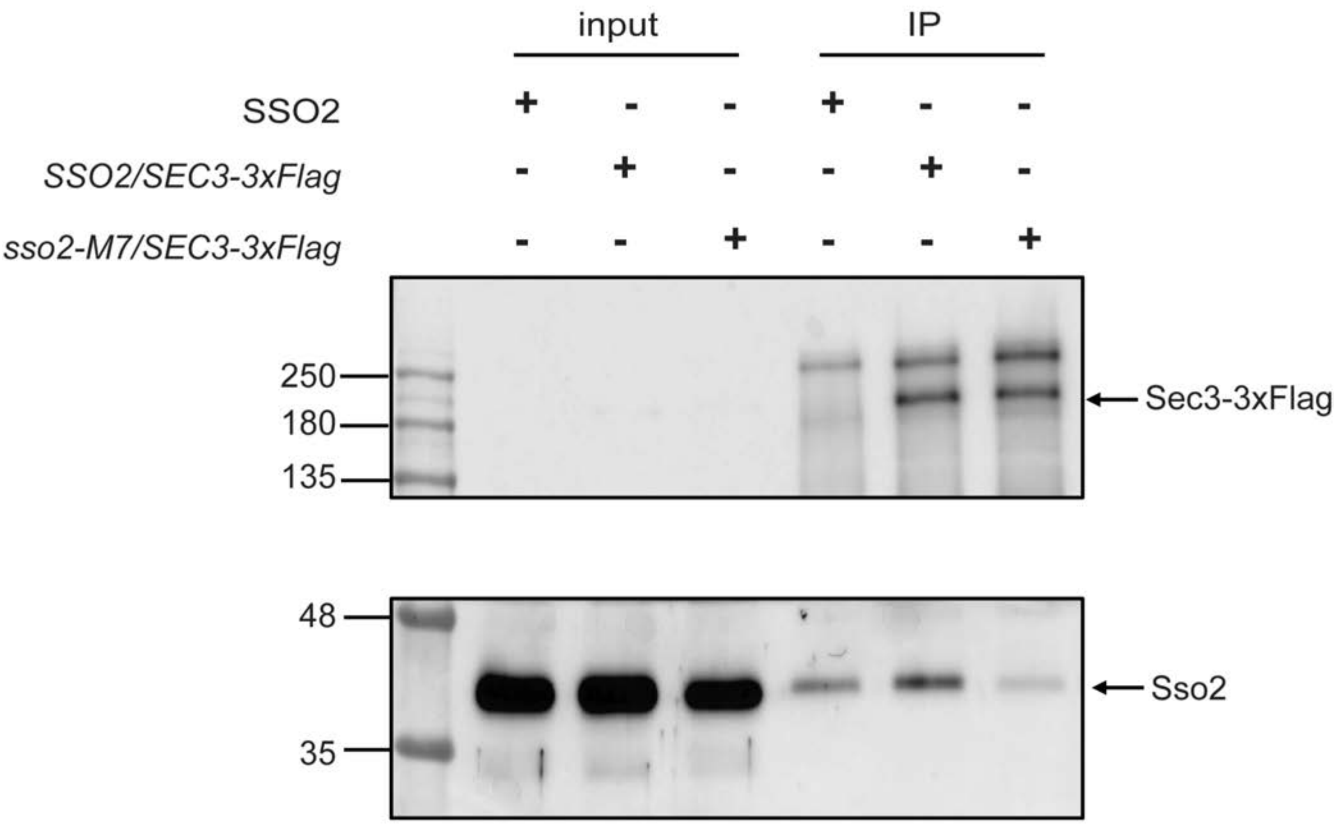
input
IP
SSO2
SSO2/SEC3-3xFlag
sso2-M7/SEC3-3xFlag
250
180
135
Sec3-3xFlag
48
35
Sso2

**Fig. S7**

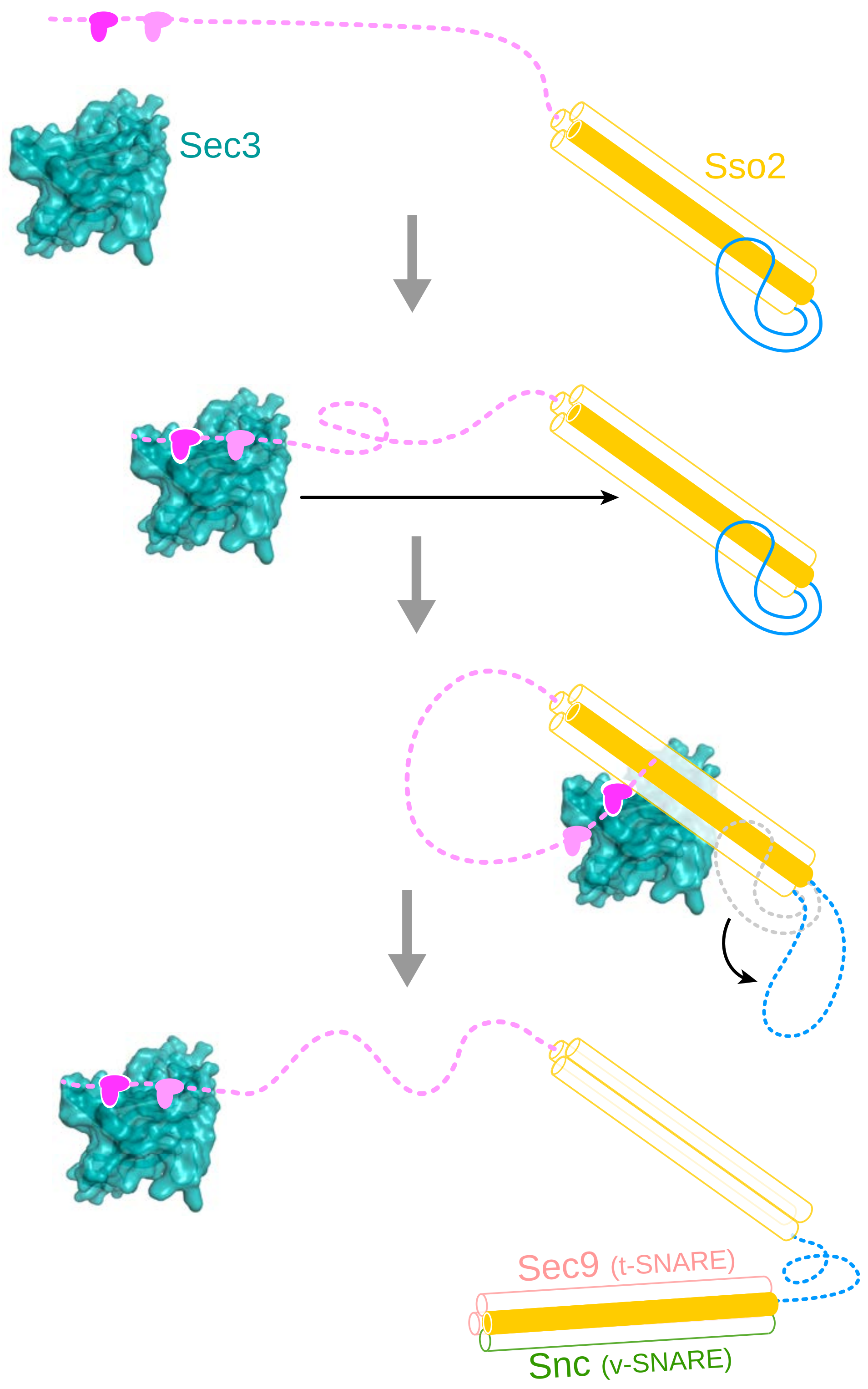

**Supplementary Table 1. Bacteria strains**

| | | |
|---|---|---|
| SFNB1223 | pRS416-TPIpro-GFP-Snc1 | Lab collection |
| NRB1303 | pRS306-Sec3-3xGFP | Lab collection |
| NRB1312 | pRS306-Sec4-GFP | Lab collection |
| NRB1644 | pRS305-Sec3-His-3xFlag (integration into SEC3 locus) | Lab collection |
| NRB1652 | hemizap-pRS306-SSO2pro-*SSO2* | This study |
| NRB1653 | hemizap-pRS306-SSO2pro-*sso2*-E8A *(sso2M1)* | This study |
| NRB1654 | hemizap-pRS306-SSO2pro-*sso2*-Y7A (*sso2M2*) | This study |
| NRB1655 | hemizap-pRS306-SSO2pro-*sso2*-N5AP6A (*sso2M3*) | This study |
| NRB1656 | hemizap-pRS306-SSO2pro-*sso2*-N5AP6AY7A (*sso2M4*) | This study |
| NRB1657 | hemizap-pRS306-SSO2pro-*sso2*-N5AP6AY7AE8A (*sso2M5*) | This study |
| NRB1658 | hemizap-pRS306-SSO2pro-*sso2*-N11AP12AY13A (*sso2M6*) | This study |
| NRB1659 | hemizap-pRS306-SSO2pro-*sso2*-N5AP6AY7AE8A-N11AP12AY13A (*sso2M7*) | This study |

**Supplementary Table 2. Yeast strains**

| | | |
|---|---|---|
| NY3305 | *Mat-a his3Δ1 lys2-Δ0 leu2Δ0 ura3Δ0 sso1Δ::KanMX4 SSO2* | This study |
| NY3306 | *Mat-a his3Δ1 lys2-Δ0 leu2Δ0 ura3Δ0 sso1Δ::KanMX4 sso2M1* | This study |
| NY3307 | *Mat-a his3Δ1 lys2-Δ0 leu2Δ0 ura3Δ0 sso1Δ::KanMX4 sso2M2* | This study |
| NY3308 | *Mat-a his3Δ1 lys2-Δ0 leu2Δ0 ura3Δ0 sso1Δ::KanMX4 sso2M3* | This study |
| NY3309 | *Mat-a his3Δ1 lys2-Δ0 leu2Δ0 ura3Δ0 sso1Δ::KanMX4 sso2M4* | This study |
| NY3310 | *Mat-a his3Δ1 lys2-Δ0 leu2Δ0 ura3Δ0 sso1Δ::KanMX4 sso2M5* | This study |
| NY3311 | *Mat-a his3Δ1 lys2-Δ0 leu2Δ0 ura3Δ0 sso1Δ::KanMX4 sso2M6* | This study |
| NY3312 | *Mat-α his3Δ1 lys2-Δ0 leu2Δ0 ura3Δ0 sso1Δ::KanMX4 sso2M7* | This study |
| NY3313 | *Mat-a his3Δ1 lys2-Δ0 leu2Δ0 ura3Δ0 sso1Δ::KanMX4 SSO2 [GFP-SNC1::URA3 CEN6]* | This study |
| NY3314 | *Mat-a his3Δ1 lys2-Δ0 leu2Δ0 ura3Δ0 sso1Δ::KanMX4 sso2M5 [GFP-SNC1::URA3 CEN6]* | This study |
| NY3315 | *Mat-a his3Δ1 lys2-Δ0 leu2Δ0 ura3Δ0 sso1Δ::KanMX4 sso-M6 [GFP-SNC1::URA3 CEN6]* | This study |
| NY3316 | *Mat-α his3Δ1 lys2-Δ0 leu2Δ0 ura3Δ0 sso1Δ::KanMX4 sso2M7 [GFP-SNC1::URA3 CEN6]* | This study |
| NY3317 | *Mat-a his3Δ1 lys2-Δ0 leu2Δ0 ura3Δ0 sso1Δ::KanMX4 SSO2[pRS306-SEC4-GFP::URA]* | This study |
| NY3318 | *Mat-a his3Δ1 lys2-Δ0 leu2Δ0 ura3Δ0 sso1Δ::KanMX4 sso2M5 [pRS306-SEC4-GFP::URA]* | This study |
| NY3319 | *Mat-α his3Δ1 lys2-Δ0 leu2Δ0 ura3Δ0 sso1Δ::KanMX4 sso2M7 [pRS306-SEC4-GFP::URA]* | This study |
| NY3320 | Mat-a his3Δ1 lys2-Δ0 leu2Δ0 ura3Δ0 sso1Δ::KanMX4 *SSO2* [pRS306-Sec3-3xGFP::URA3] | This study |
| NY3321 | Mat-a his3Δ1 lys2-Δ0 leu2Δ0 ura3Δ0 sso1Δ::KanMX4 *sso2M5* [pRS306-Sec3-3xGFP::URA3] | This study |
| NY3322 | Mat-α his3Δ1 lys2-Δ0 leu2Δ0 ura3Δ0 sso1Δ::KanMX4 *sso2M7* [pRS306-Sec3-3xGFP::URA3] | This study |
| HY3447 | *Mat-a his3Δ1 lys2-Δ0 leu2Δ0 ura3Δ0 sso1Δ::KanMX4 SSO2* [pRS305-Sec3-His-3xFlag | This Study |
| HY3448 | *Mat-α his3Δ1 lys2-Δ0 leu2Δ0 ura3Δ0 sso1Δ::KanMX4 sso2M7* [pRS305-Sec3-His-3xFlag] | This study |
